# Minimally-overlapping words for sequence similarity search

**DOI:** 10.1101/2020.07.24.220616

**Authors:** Martin C. Frith, Laurent Noé, Gregory Kucherov

## Abstract

Analysis of genetic sequences is usually based on finding similar parts of sequences, e.g. DNA reads and/or genomes. For big data, this is typically done via “seeds”: simple similarities (e.g. exact matches) that can be found quickly. For huge data, sparse seeding is useful, where we only consider seeds at a subset of positions in a sequence.

Here we study a simple sparse-seeding method: using seeds at positions of certain “words” (e.g. `ac, at, gc`, or `gt`). Sensitivity is maximized by using words with minimal overlaps. That is because, in a random sequence, minimally-overlapping words are anti-clumped. We provide evidence that this is often superior to acclaimed “minimizer” sparse-seeding methods. Our approach can be unified with design of inexact (spaced and subset) seeds, further boosting sensitivity. Thus, we present a promising approach to sequence similarity search, with open questions on how to optimize it.

## 1 Introduction

### 1.1 Seeds

Finding similar sequences, in large data, is typically done via “seeds”: simple similarities that can be found quickly. The simplest type of seed is exact matches of a given length, e.g. 10 letters for DNA. The seed length affects the sensitivity and run time: shorter seeds are more sensitive, but find more hits that must then be checked. By lengthening the seeds, we can arbitrarily reduce the run time of the downstream steps, but not the time and memory usage for finding the seeds.

### 1.2 Sparse seeds

An alternative way to reduce time and/or memory use is sparse seeding. The simplest way is to only use seeds starting at every *n*th position in one of the two sequences being compared.

Note that, if we only use seeds at every *n*th position in *both* sequences, the sensitivity will be poor. E.g. if there are long similar segments, without insertions or deletions, starting at coordinate *x* in the first sequence and coordinate *x* + 1 in the second sequence, they are never hit if *n* > 1.

Sparse seeding reduces sensitivity, but we could then increase the sensitivity by shortening the seeds. This raises the prospect of reducing run time and/or memory use without loss of sensitivity.

### 1.3 Sparsity via words

An intriguing idea is to achieve sparsity by selecting seeds starting at positions of certain words. For example, if we only use seeds starting with a (Paul Horton, personal communication), we achieve 4-fold sparsity in both sequences without huge loss of sensitivity. We can imagine more complex variants, e.g. use seeds starting with any of these words: ac, at, gc, gt. Surprisingly, it makes a difference: it is better to use words that have minimal chance of overlapping. That is because, in a random sequence, minimally-overlapping words occur with more uniform spacing, i.e. they are anti-clumped or under-dispersed; equivalently, their number of occurrences has lower variance.

### 1.4 Minimizers

A related idea is minimizers [SWA03, RHH^+^04]. This method only uses seeds starting at positions that are “minimum” in any window of *w* consecutive positions (e.g. *w* = 7). Various definitions of “minimum” can be used; the simplest is: the sequence (i.e. suffix) starting at this position is alphabetically minimum. This is somewhat like using seeds starting with `a`. More complex orderings can be used, e.g. compare two suffixes using order `c<a<t<g` at odd-numbered bases and `g<t<a<c` at even-numbered bases, so that `cgcg`… is the minimum possible suffix [RHH^+^04].

The resulting degree of sparsity is not obvious, and it depends on the ordering [MPB^+^17]. Typically, a fraction 2/(*w* + 1) of positions is selected [SWA03].

Another related idea is universal *k*-mer hitting sets [OPM^+^17]. This means a set of length-*k* words, such that every possible length-*L* sequence contains at least one of the words. Recent studies have defined minimizer orderings based on universal *k*-mer hitting sets, resulting in high sparsity for a given *w* [OPM^+^17, MPB^+^17, MDK18].

Minimizers have been described as “a central recent paradigm” [OPM^+^17]: they have been widely used for sparse seeding (e.g. [Li18, JKD^+^18]) and other applications (e.g. [LKH^+^13, WS14, DKGDG15]).

### 1.5 Spaced and subset seeds

So far we have considered exact-match seeds, but inexact seeds are also used. One variant is spaced seeds, which allow mismatches at some fixed positions in the seed (e.g. positions 3 and 5 out of 9). Spaced seeds are often superior to exact seeds [MTL02], because their hits are less concentrated in overlapping clumps. Thus, spaced seeds have been designed by minimizing their “overlap complexity” [II07], which is similar to minimizing the variance in number of hits [HLO^+^16].

Subset seeds are a further generalization: they allow *some* mismatches (e.g. `a↔g` and `c↔t`) at fixed positions [NK04]. This is useful for DNA, because `a↔g` and `c↔t` substitutions (termed “transitions”) are often more frequent than the other types of substitution (“transversions”). Transition seeds have also been designed for use with every-*n*th sparsity [FN14].

### 1.6 Repeats

Natural DNA has many repeats, which are the main difficulty for similarity search. For example, a primate genome may have a million Alu elements, so naive comparison of such genomes yields an unmanageable 10^12^ significant similarities. Our practical aim cannot be to find all significant similarities, but rather orthologs and/or strongest similarities. In any case, a seeding method must avoid getting too many repetitive seeds. One solution is to omit high-frequency seeds, another is to use variable-length seeds that are made longer until they are sufficiently rare [Csű04, KWS^+^11].

### 1.7 Non-overlapping words

Since we are interested in minimally-overlapping words, let us consider non-overlapping words. A basic question is: what is the maximum possible number of non-overlapping words of some length *k*? That is, given an alphabet of size *a* (so there are *a^k^* possible words), what is the maximum possible number of words where no proper prefix of any word equals a proper suffix of any word? This seems hard to answer in general [Bla15].

The following construction has been suggested for getting a large number of non-overlapping words [Bla15]. Divide the alphabet into two subsets, e.g. {a} and {c, *g,t}*, and choose a prefix length *j* (0 < *j* < *k*). These words have no overlaps: words whose first *j* letters are from the 1st subset, whose (*j* + 1)th and *k*th letters are from the 2nd subset, and whose letters between *j* + 1 and *k* have no run of ≥ *j* letters from the 1st subset.

## 2 Methods

The methods are available at: https://gitlab.com/mcfrith/noverlap

### 2.1 Mean and variance

Given a set of length-*k* words, let us consider their occurrence in a random i.i.d. length-s sequence. The expected number of total occurrences is:

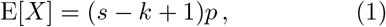

where *p* is the total probability of any of the words occurring at a given position. The variance in occurrence number is:

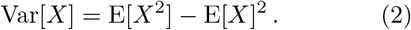

To calculate this, let us define index variables *I^j^* as 1 if any of the words occurs at position *j*, else 0. So:

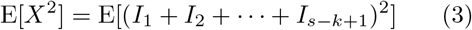

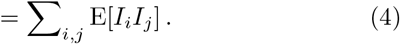

If we define *l* = <*i − j*<, then

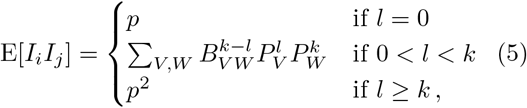

where 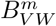 is defined to be 1 if the length-*m* suffix of word *V* equals the length-*m* prefix of word *W*, else 0. Also, 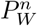 is the product of probabilities of the first *n* letters in word W. Thus, assuming that *s* ≥ 2*k* − 2 (see the Supplement):

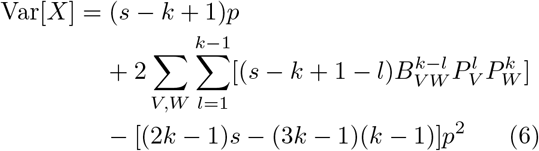

For circular sequences, the formulas are simpler (assuming *s* ≥ 2*k* − 1):

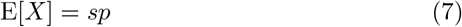

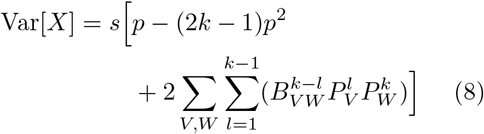

These formulas also apply to linear sequences when *s ∫ k*. With these formulas, the variance-to-mean ratio, also called index of dispersion, is independent of the sequence length. The formulas also simplify for linear sequences with *s* = 2*k* − 1 (the smallest s where all kinds of pairwise overlap contribute):

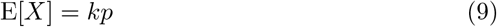

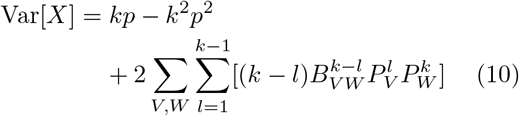

### 2.2 Simulated sequences

To test homology detection, DNA sequences were simulated with the T92 model of evolution [Tam92]. This model has three input parameters: `gc-content`, transition/transversion rate ratio *κ*, and PAM substitution distance (Table 1).

**Table 1:** Parameters of the T92 DNA model

| PAM | $\kappa$ | %g+c | %identity | transitions per transversion |
| --- | --- | --- | --- | --- |
| 20 | 1 | 50 | 82 | 0.5 |
| 50 | 1 | 50 | 64 | 0.5 |
| 20 | 3 | 50 | 83 | 1.4 |

For each test, 100 000 pairs of DNA sequences were simulated. The default parameters, unless specified otherwise, are: %g+c = 50, *κ* = 1 (unbiased), PAM = 20, sequence length = 100. A seeding method was deemed to find a pair of sequences if it found at least one match at identical coordinates of the pair.

To test specificity, two unrelated length-10^6^ sequences were generated, and the number of seed pair matches counted. This is a proxy for the computational cost of checking all the seed hits.

## 3 Results

### 3.1 Non-overlapping DNA words

The maximum possible number of non-overlapping DNA words, for word length *k* = 2–6 (Table 2), was found by brute force clique search [KJ07]. For *k* < 6, Blackburn’s construction [Bla15] achieves this maximum. For *k* = 2, a maximum set is `ry` (`r = a` or `g, y = c` or `t`). In general, `abb`… (`b` = any base except `a`) is a good way to get non-overlapping words, and a nice generalization of Horton’s idea.

**Table 2:** Non-overlapping DNA words. `r` = {`a, g`}; `y` = {`c, t`}; `b` = {`c, g, t`}.

| word length | constructed words | number | maximum number |
| --- | --- | --- | --- |
| 2 | ry | 4 | 4 |
| 3 | abb | 9 | 9 |
| 4 | abbb | 27 | 27 |
| 5 | abbbb | 81 | 81 |
| 6 | abbbbb | 243 | 251 |

### 3.2 Every *n*th sparsity

We first tested every-*n*th sparsity (only using seeds starting at every *n*th position in one of the two sequences being compared), with exact-match seeds. We defined “sensitivity” as % of sequence pairs with ≥ 1 seed match at homologous positions. As expected, if we increase sparsity without changing the seed length, both sensitivity and random hit count decrease (Figure 1). If we then shorten the seeds, the sensitivity and random hit count increase. The important result is that higher sparsity has lower sensitivity *for a given random hit count*. The exception is *n* = 2, which is no worse than *n* = 1, indeed giving us something for nothing: sparsity at no cost.

**Figure 1:**
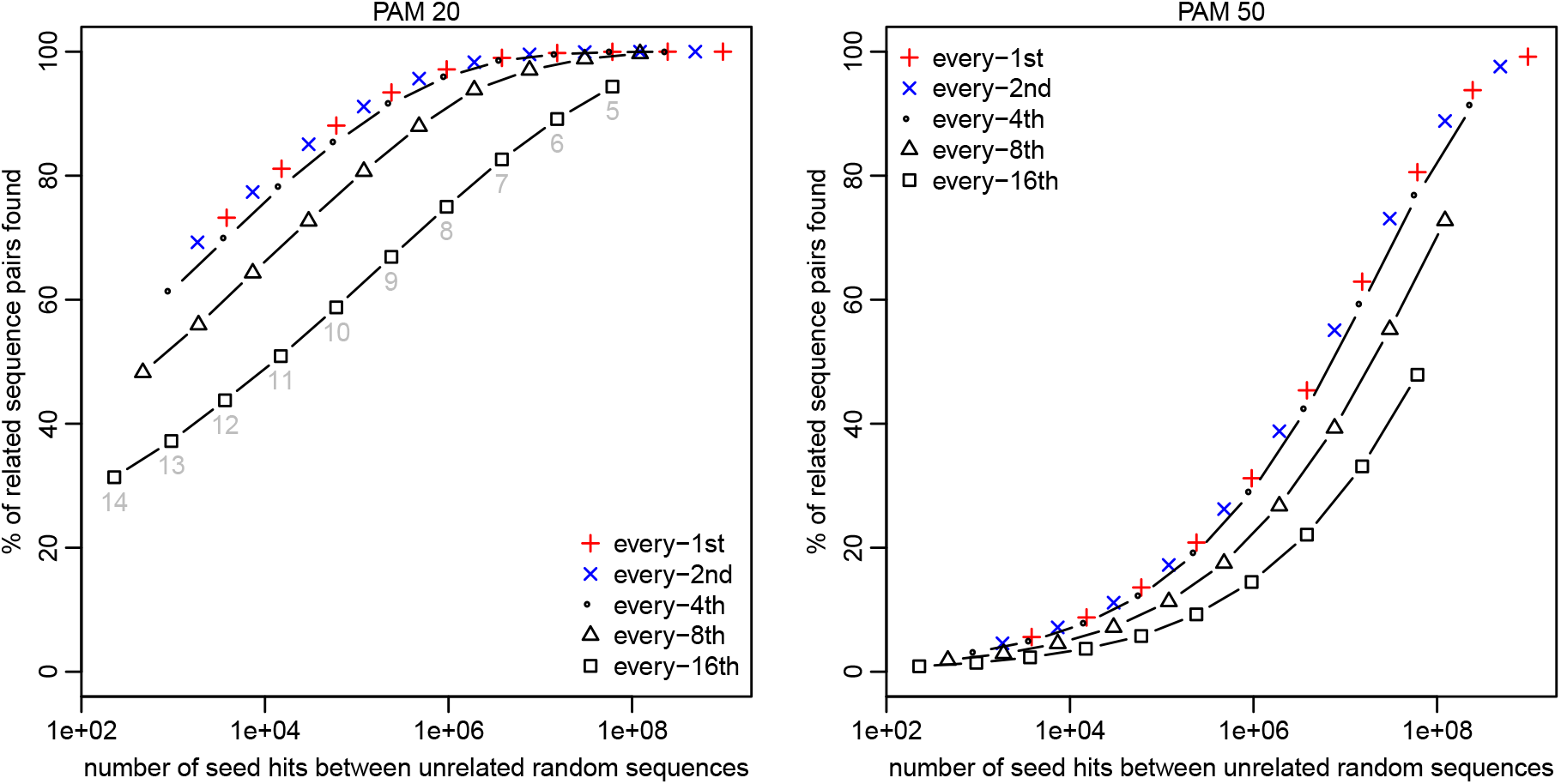
Sensitivity (y-axis) and spurious hit count (x-axis) for exact-match seeds with every-*n*th sparsity. Sensitivity was measured on sequence pairs with PAM distance 20 (left panel) or 50 (right panel). Seed lengths 5–14 were tested, shown in gray in the left panel.

A plausible explanation for why *n* = 1 is not better than *n* = 2 is that highly-overlapping seeds provide little independent information. This is also why spaced seeds are better than exact-match seeds. Thus, it would be interesting to compare *n* = 1 to *n* =2 using optimized spaced/subset seed patterns: this was done previously, and *n* = 2 was worse [FN14].

### 3.3 Sparsity via words

Let us now see how seeds starting with a compare to every-4th seeding. For a given seed length, the random hit counts are the same (as expected), but seeds starting with a have lower sensitivity (Figure 2A). This is not too surprising, because every-4th seeding is sparse in just one sequence, but seeds starting with a are sparse in both sequences. Seeds starting with `ry` also have the same random hit counts, and their sensitivity is closer to (but still less than) that of every-4th seeds. On the other hand, seeds starting with `rr` have worse sensitivity. This supports the idea that non-overlapping words are good and highly-overlapping words are bad.

**Figure 2:**
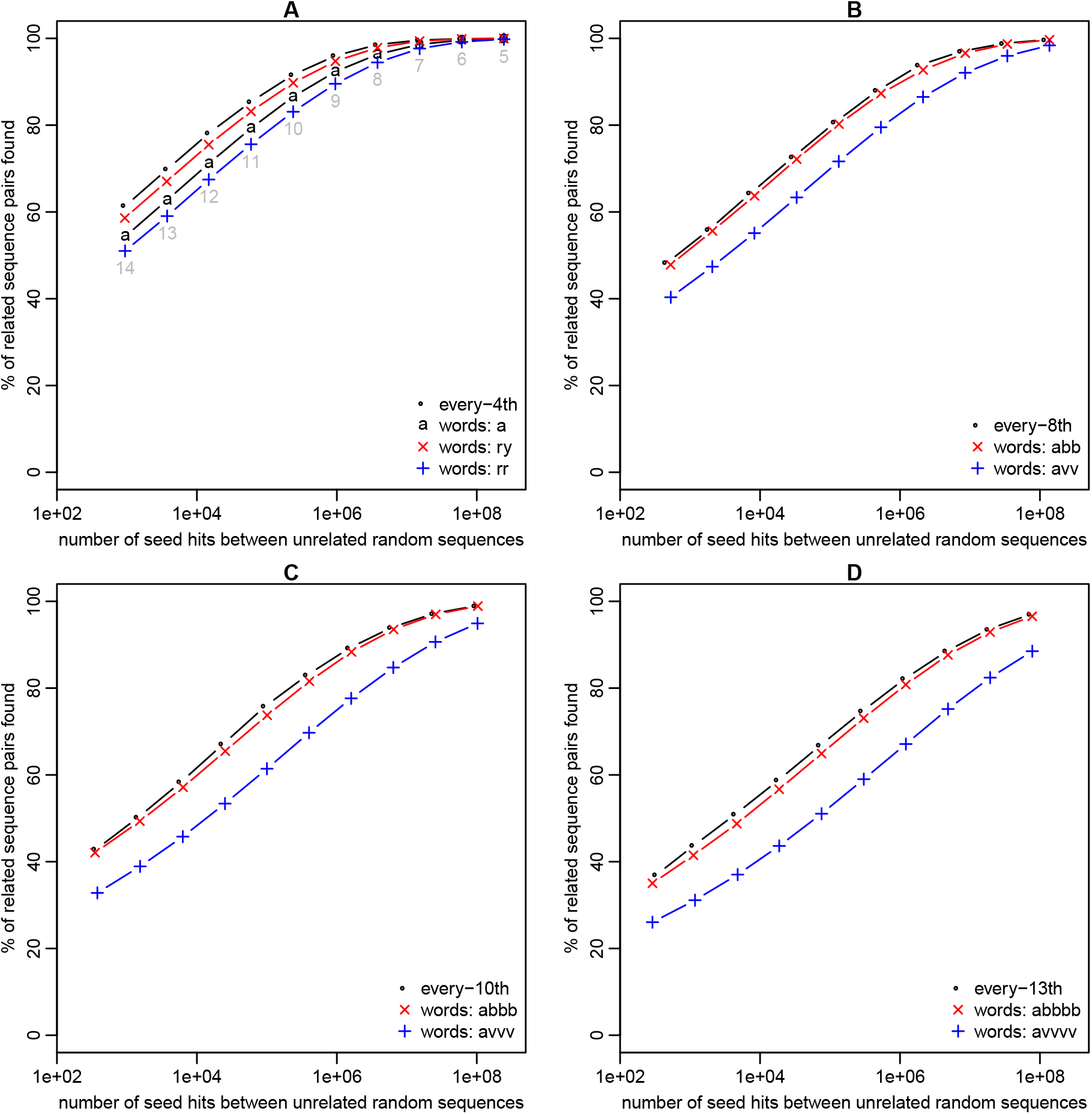
Sensitivity (y-axis) and spurious hit count (x-axis) for exact-match seeds with every-*n*th or word-based sparsity. Sensitivity was measured on sequence pairs with PAM distance 20. Seed lengths 5–14 were tested, as shown in panel **A**.

Seeds starting with abb have a sparsity factor of 4^3^/3^2^ ≈ 7.1, and they perform slightly worse than every-8th seeds (Figure 2B). On the other hand, they perform better than seeds starting with `avv` (`v` = any base except `t`). Seeds starting with `abbb` (sparsity 9.5) or `abbbb` (sparsity 12.6) show similar results (Figure 2C–D), confirming the advantage of non-overlapping words.

### 3.4 Minimal-variance words

We can perhaps do better by using longer words with some overlap. Seeds starting with `ry` are the same as seeds starting with `ryn` (where `n` is any base), so it may be better to replace `ryn` with a less-overlapping set of length-3 words.

It is not obvious how best to quantify “amount of overlap”, but one idea is to use variance of occurrence number in random sequences. Let us try these two measures of overlap: VMR1 (variance-to-mean ratio from Equations 7,8) and VMR2 (from Equations 9,10).

It is also unclear how to find a set of words that minimizes VMR1 or VMR2, because the number of possible sets is enormous. Brute-force search is feasible if we restrict ourselves to a 2-letter `ry` alphabet.

Such words can indeed boost sensitivity. For example, the words `rrry,ryrr,ryyr,yyyr` have lower VMR2 than `rynn` (Table 3), and seeds starting at these words have better sensitivity (Figure 3A). We can do better still with eight length-5 words (Table 3, Figure 3A).

**Table 3:**
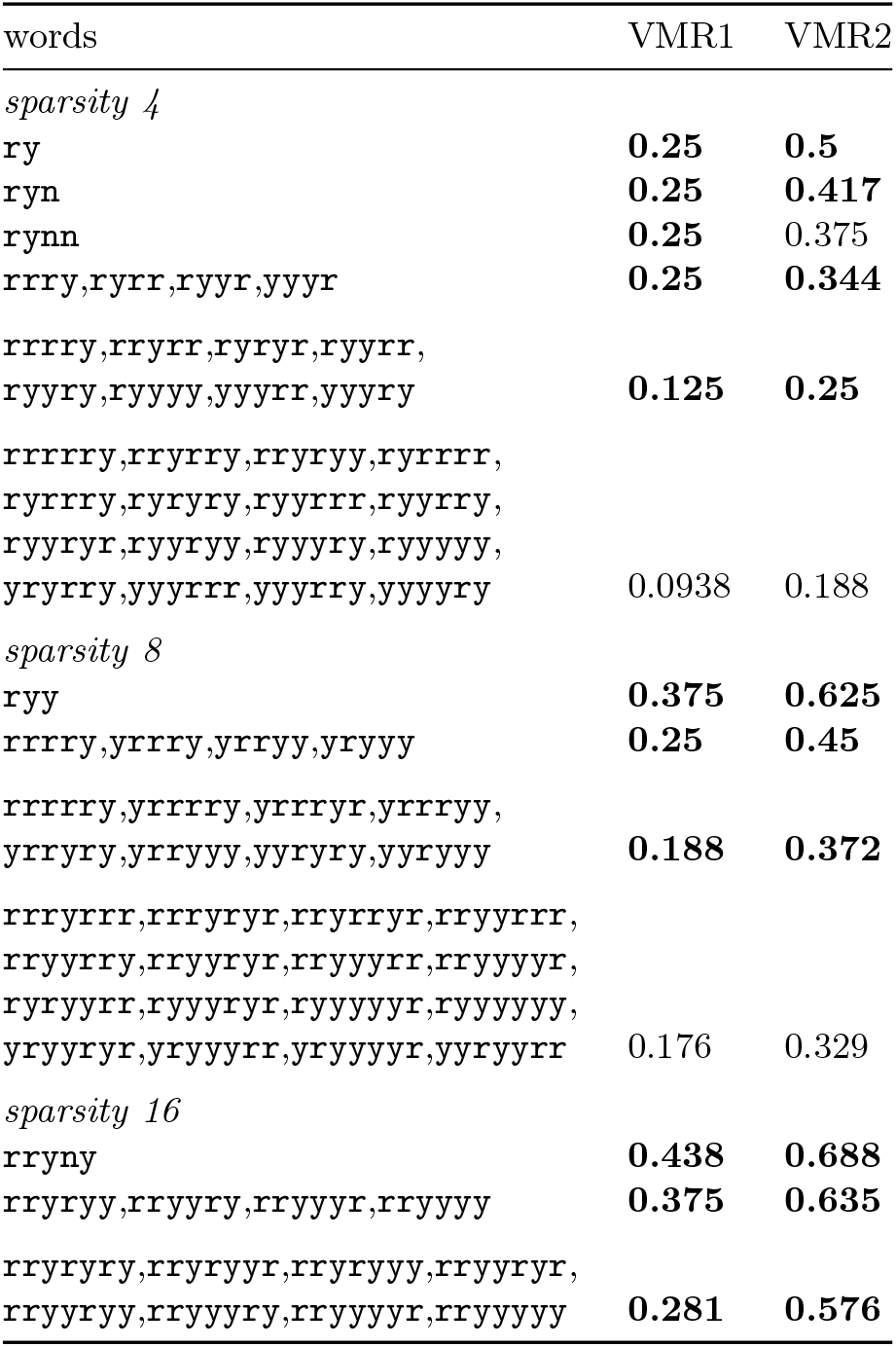
Variance-to-mean ratios. Bold values are known to be the minimum possible, for that sparsity and word length.

**Figure 3:**
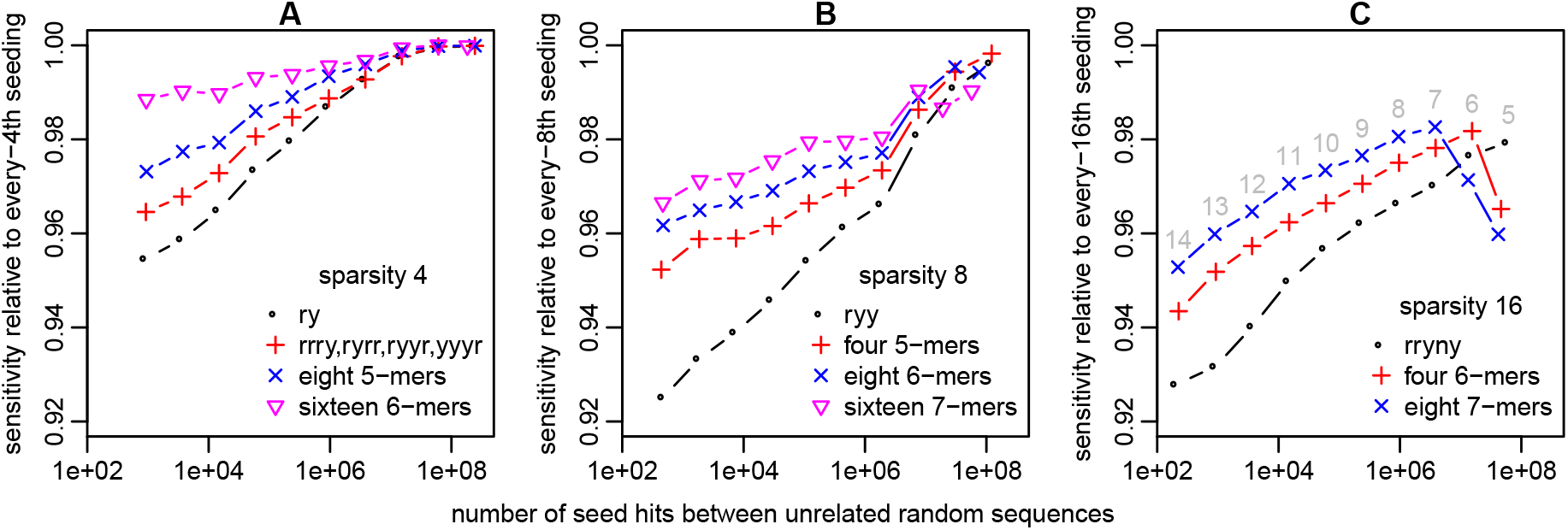
Sensitivity (y-axis) and spurious hit count (x-axis) for exact-match seeds with word-based sparsity. Sensitivity was measured on sequence pairs with PAM distance 20. Seed lengths 5–14 were tested, as shown in panel **C**. In this figure, the sensitivity is shown relative to every-*n*th sparsity: (% of related sequence pairs found by word-restricted seeds) / (% of related sequence pairs found by every-*n*th seeds).

For even longer words, our brute-force search was too slow, so we switched to a heuristic search method (simulated annealing) that does not guarantee to find the minimum possible VMR. This found a set of sixteen length-6 words (Table 3), with even better sensitivity (Figure 3A).

On the other hand, we also found cases where words with lower VMR1 and VMR2 have worse sensitivity (see the Supplement). Thus, a better criterion for choosing a set of words is still to be designed.

Figure 3B–C shows other examples where longer minimum-variance words (from Table 3) improve sensitivity, with no change in sparsity or random hit rate. However, the longer words perform badly when they exceed the seed length (e.g. blue and red points at the right of Figure 3C), showing a danger of too-long words.

## 3.5 Minimizers

We next tested minimizers, with three orderings:

- Alphabetic order.
- `cg`-order, where `cgcg`… is the minimum sequence. This is representative of methods that have been used in practice [MPB^+^17].
- `abb`-order. This is a novel ordering, inspired by non-overlapping `abb`… words. It compares two suffixes using order `a<c<g<t` at the first position and `t=g=c<a` at all subsequent positions.

Let us first see the sparsity (average distance between seed start coordinates) of these orderings. Alphabetic minimizers have the lowest sparsity (highest density) for a given window length *w*, and `cg` minimizers have higher sparsity (Figure 4), as reported previously [MPB^+^17]. Interestingly, abb minimizers have even higher sparsity for *w* > 10.

**Figure 4:**
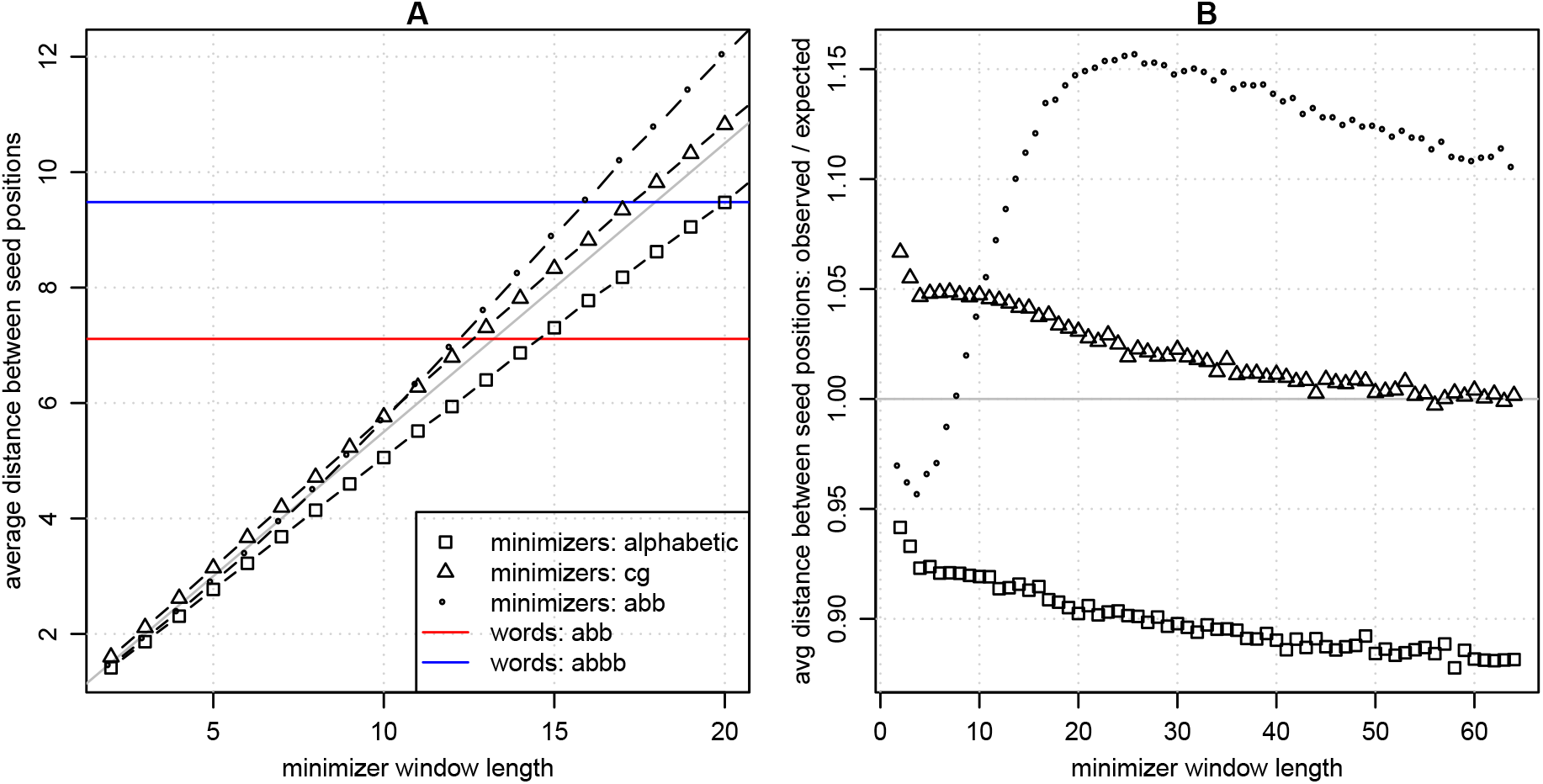
Sparsity of minimizers, with three orderings. The diagonal gray line in **A**, and the horizontal gray line in **B**, show the expected minimizer sparsity (*w* + 1)/2.

Now let us see the sensitivity of these minimizers. Taking alphabetic minimizers as an example, if we increase the window size *w* without changing the seed length, the sensitivity and random hit rate both decrease (Figure 5), as expected. If we then shorten the seeds, the sensitivity and random hit rate increase. Overall, higher *w* results in lower sensitivity for a given random hit count.

**Figure 5:**
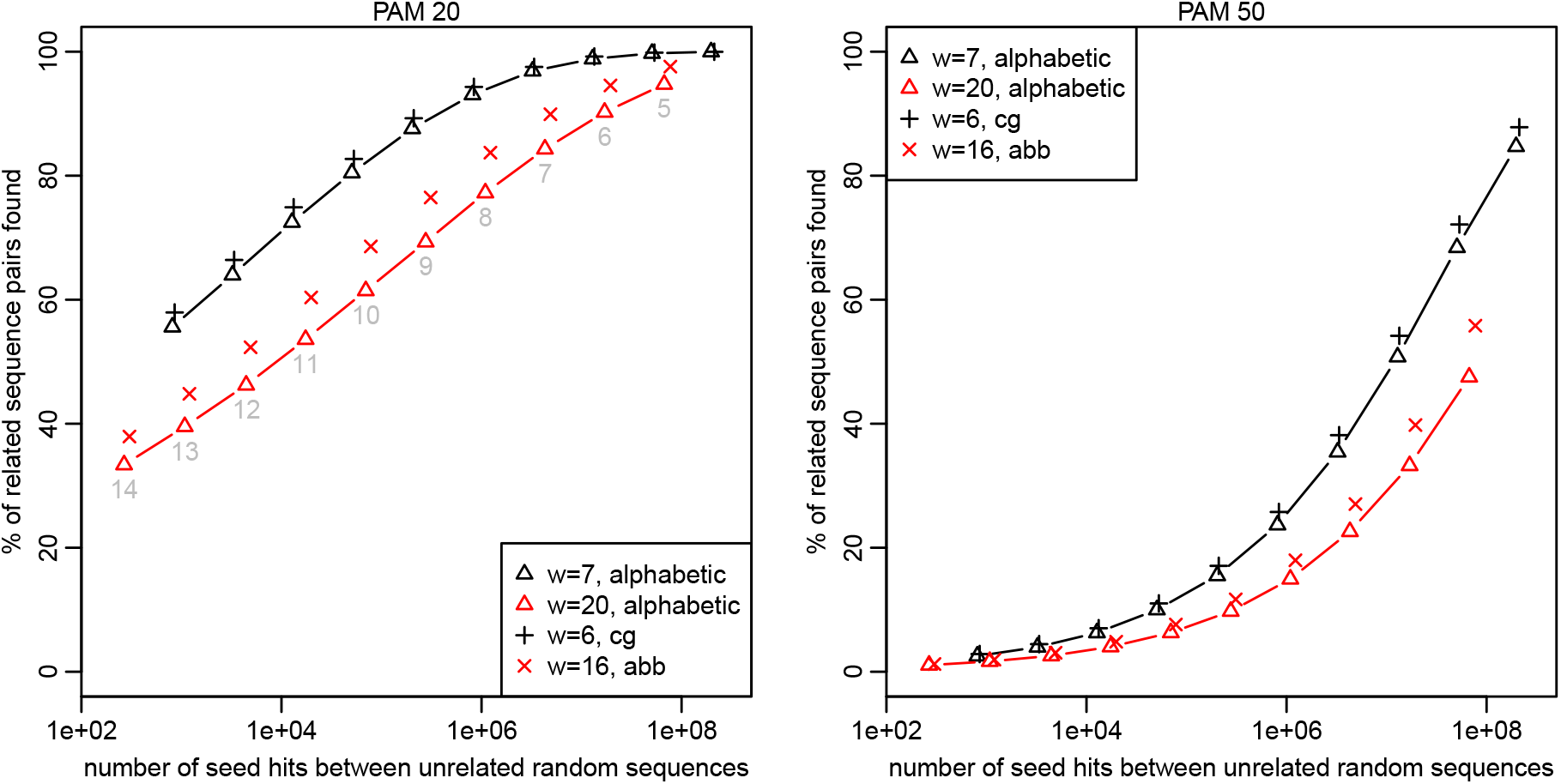
Sensitivity (y-axis) and spurious hit count (x-axis) for exact-match seeds at minimizer positions. “w” means window length. Seed lengths 5–14 were tested, shown in gray in the left panel.

To fairly compare the three kinds of minimizer, we should compare them using *different* window sizes that achieve the *same* sparsity. Based on Figure 4A, alphabetic minimizers with *w* = 7 are comparable to `cg` minimizers with *w* = 6, and alphabetic minimizers with *w* = 20 are comparable to `abb` minimizers with *w* = 16. Comparing them thus, `cg` and `abb` minimizers are better than alphabetic minimizers (Figure 5). This supports the idea that higher sparsity for a given *w* improves homology search, which does not seem to have been clearly shown before.

## 3.6 Minimizers versus words

To fairly compare minimizers with words, we should use minimizer window sizes that produce the same sparsity as the words. Figure 6A,B compares words to minimizers with slightly lower sparsity (higher density), giving an unfair advantage to the minimizers. Seeds starting with a perform worse than alphabetic minimizers for PAM distance 20 (Figure 6A), but better for PAM distance 50 (Figure 6B). On the other hand, seeds starting at non-overlapping (`ry`) or minimum-variance words perform better than alphabetic or `cg` minimizers, at both PAM distances.

**Figure 6:**
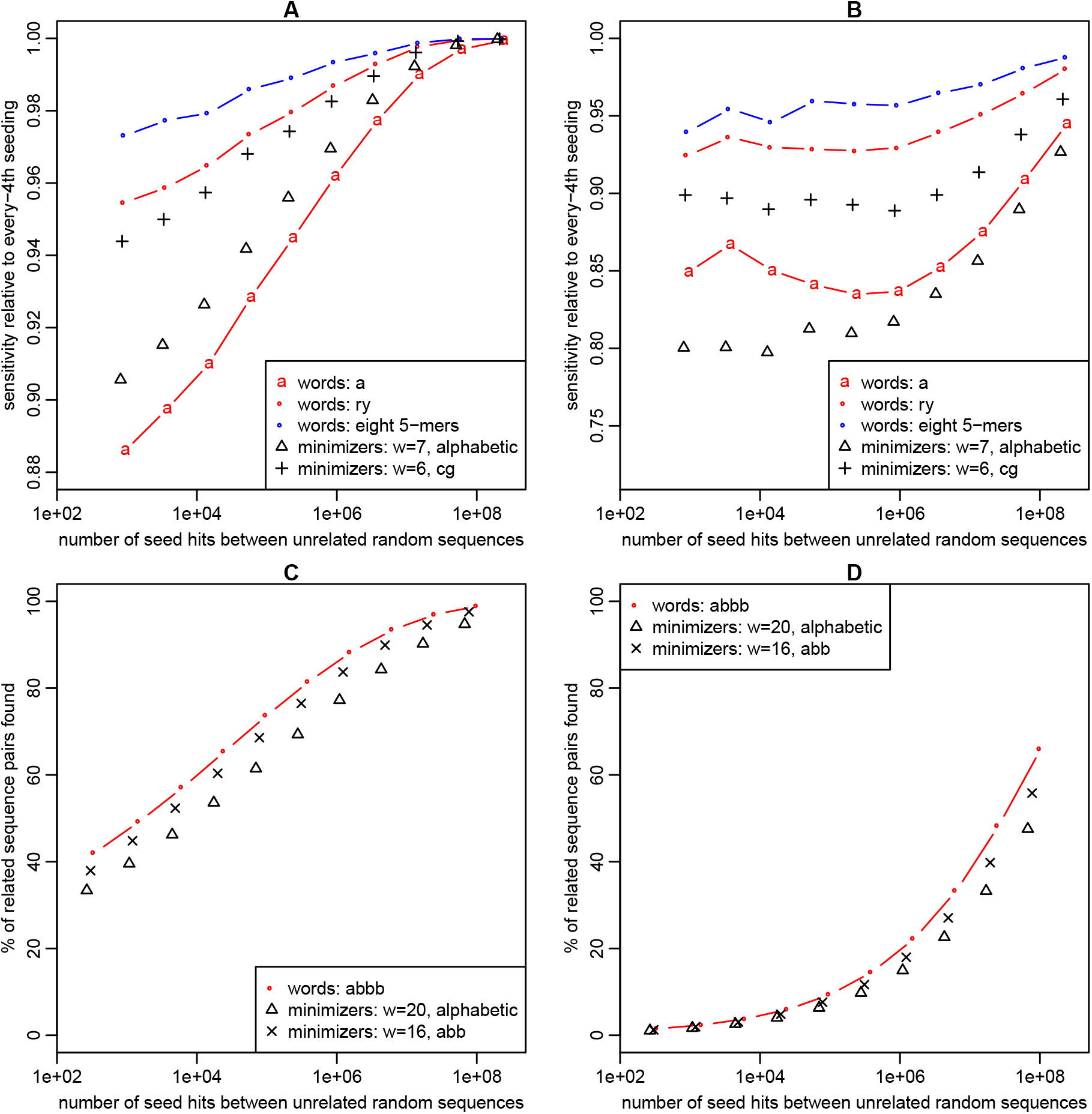
Sensitivity (y-axis) and spurious hit count (x-axis) for exact-match seeds at either word positions or minimizer positions. Seed lengths 5–14 were tested. Sensitivity was measured on sequence pairs with PAM distance 20 (**A, C**) or 50 (**B, D**).

Figure 6C,D compares words to minimizers with about the same sparsity. Seeds at positions of `abbb` perform slightly better than alphabetic or `abb` minimizers. In more detail: for a given seed length, the minimizers have worse sensitivity but slightly better specificity.

Next, we compared word-based seeding to the minimizer scheme of the minimap2 software [Li18]. This scheme uses exact-match seeds of length 15, with minimizer window *w* = 10, and an ordering from a particular hash function applied to each 15-mer. The expected density is 2/(*w* + 1) = 0:1818, but we empirically found a slightly higher density, 0.185{0.188, in both random and real sequences. We compared this to twelve length-6 `ry` words (density 12=26 = 0:1875) that minimize VMR2: `rrrrry, rryrrr, rryrry, rryyrr, rryyry, ryryrr, ryyyrr, ryyyry, ryyyyr, yryyrr, yryyry, yyyyyr`.

For this test, random fragments of size 1000 were drawn from human (GRCh38) chromosome 22, then mutated by the PAM process, and the number of conserved 15-mer seeds was counted. At PAM distance 0, minimap has more seeds, i.e. higher density (Figure 7). Nevertheless, at PAM distance ≥ 1, minimap has fewer conserved seeds.

**Figure 7:**
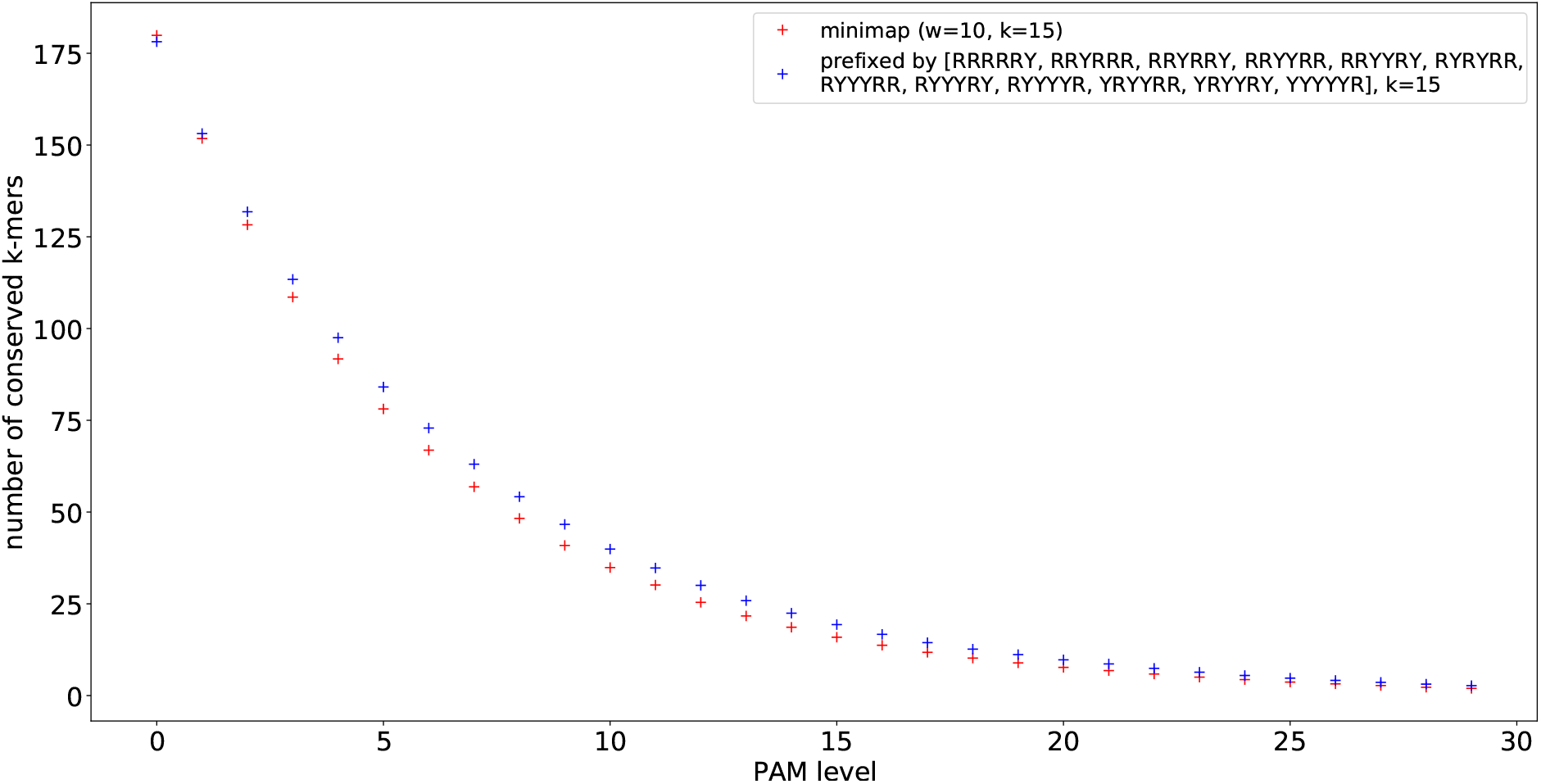
Sensitivity (y-axis) at different evolutionary distances (x-axis), for minimap seeds and word-based seeds. Here, “sensitivity” is the average number of conserved seeds over 1000 pairs of length-1000 sequences from human chromosome 22.

**Figure 8:**
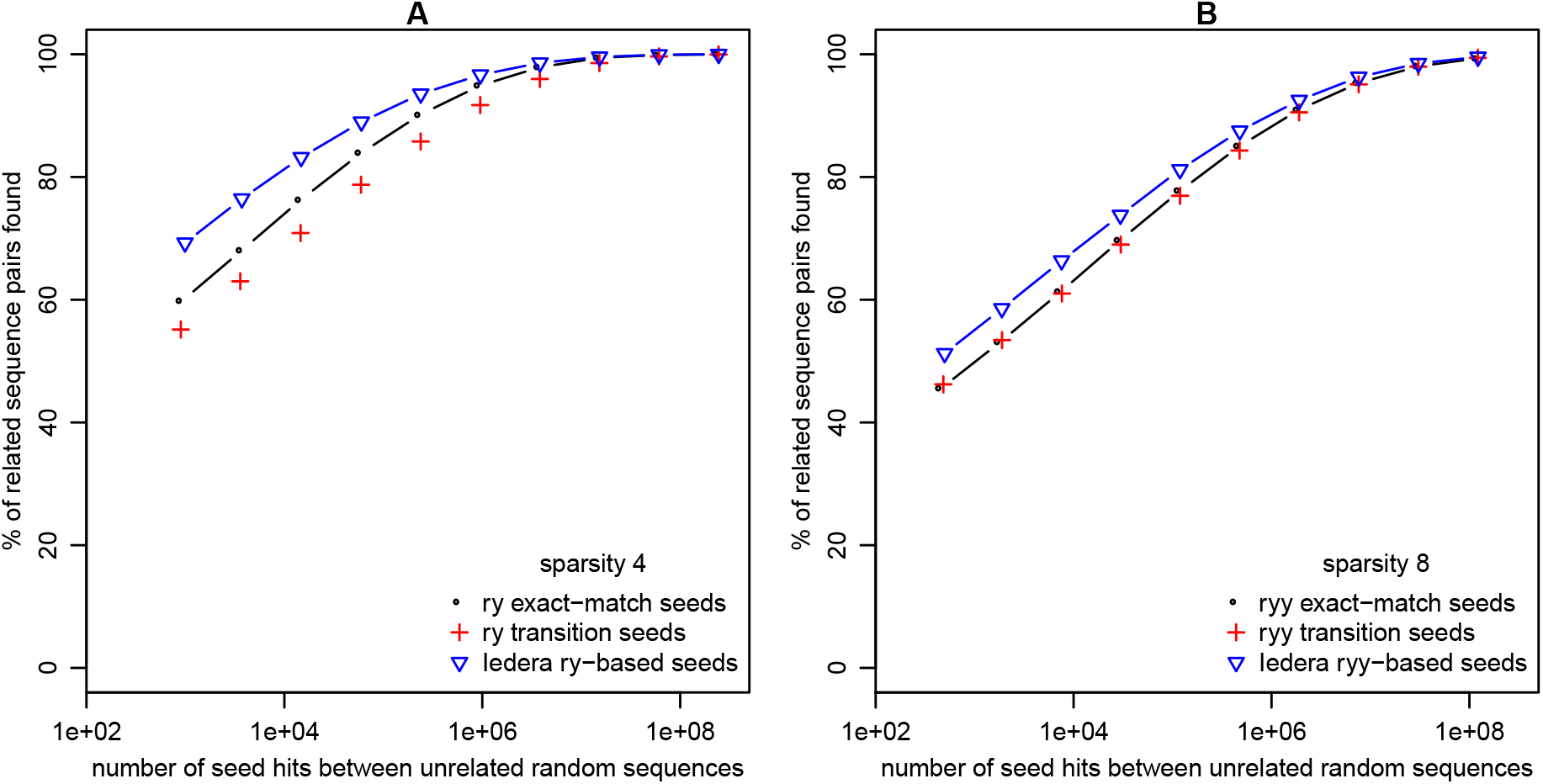
Sensitivity (y-axis) and random hit count (x-axis) of seeding methods, for sequences with transition/transversion bias (*κ* = 3) and PAM distance 20. Seed weights 5–14 were tested. “Transition seeds” allow transition substitutions at all positions.

## 3.7 Unification with subset seeds

A straightforward generalization of subset seeds incorporates word-restricted seeding. Recall that subset seeds allow some mismatches (e.g. `a↔g` and `c↔t`) at fixed positions. More generally, we can allow any subset of the 16 possible types of match and mismatch.

An example of such a generalized subset seed pattern is: ANNRYrn@y. This specifies seeds of length 9. Positions with `A` (in this case, the 1st position) allow `a:a` matches only. Positions with `N` allow any match. Positions with `n` allow any match or mismatch. Positions with `R` allow purine matches only: `a:a` or `g:g`. Positions with `r` allow purine matches or mismatches: a:a, g:g, a:g, g:a. Positions with `Y` or `y` likewise allow pyrimidines (`c` and `t`). Finally, positions with @ allow any match or transition.

Such a seed pattern has two important properties: sparsity and weight. Sparsity means rarity of compatible positions in a single random sequence. Weight indicates unlikelihood of a chance match to a compatible position: for example, weight 5 means the same unlikelihood as a length-5 exact match.

The hard problem is to design good seed patterns for finding sequences related by a given PAM distance, transition/transversion bias, etc. Fortunately, the seed design software Iedera already allowed this kind of generalized subset seed [KNR06]. Here, we used it to design seeds for PAM=20 and *κ*=3. To constrain the search space in this preliminary study, we only considered patterns based on `ry`, i.e. having one `R` or `r` and one `Y` or `y`, and likewise patterns based on `ryy`. The only other pattern symbols allowed were N, @, and up to 5 ns. Up to 10 transition-tolerant positions other than `n` were allowed. The resulting patterns are in Table 4. Many other patterns are equally good; we broke ties by preferring ones that start with `RY` or `RYY`.

One notable result is that `n` positions are useful in the `ry`-based seeds, but not the `ryy`-based seeds. This is presumably because n positions make overlapping seeds more independent, but sparser seeds have fewer overlaps.

**Table 4:**
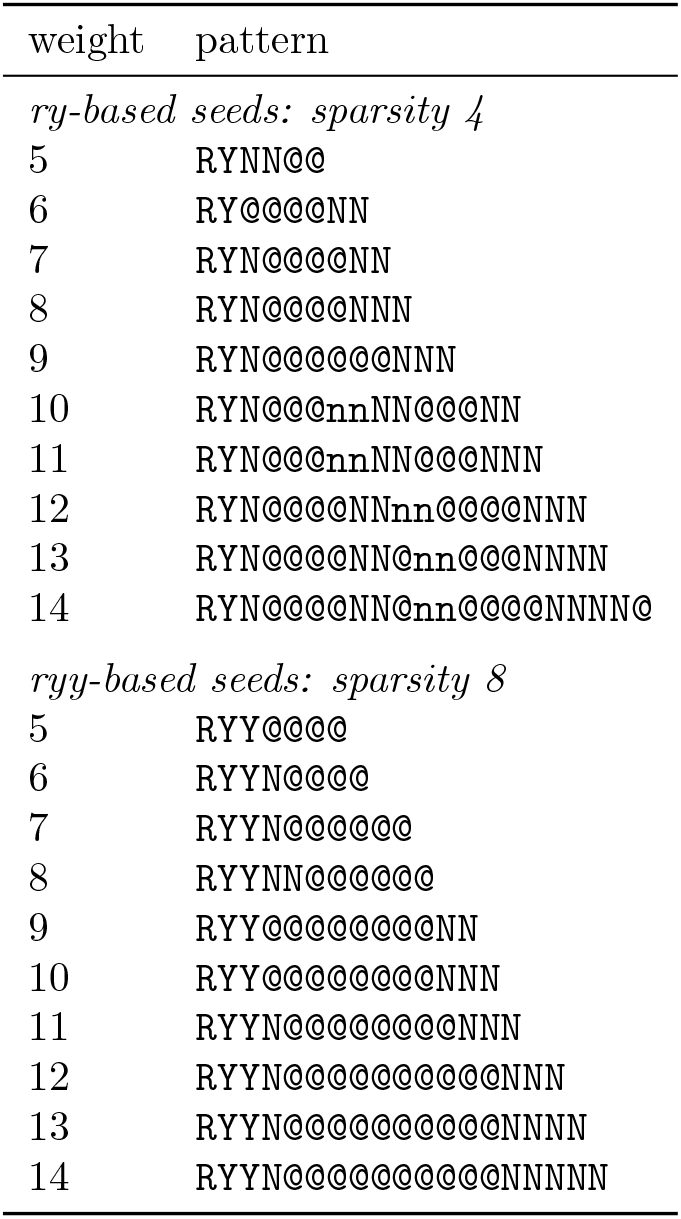
Seed patterns designed by Iedera for PAM 20, *κ* = 3, alignment length 64.

## 4 Discussion

### 4.1 When to use sparsity

The aim of seeding methods is to maximize sensitivity while minimizing computational cost (time and memory). Computational cost has two parts: the cost of finding seed matches (*c*^1^) and the cost of processing them (*c*^2^). Sparsity need not reduce sensitivity, if the seeds are shortened, but it usually increases random seed hits (i.e. *c*^2^) for a given sensitivity (Figure 1). A notable exception is exactmatch seeds and every-*n*th sparsity with small *n* (e.g. *n* = 2), which does not increase random hits for a given sensitivity (Figure 1). Typically, however, sparsity is beneficial only when long (or rare) seeds do not sufficiently reduce the computational cost.

### 4.2 When to use every-*n*th sparsity

Every-*n*th sparsity has better sensitivity per random hits (*c*^2^) than either minimizers or word-restricted seeds, see also [AT18]. So it should be preferred unless its *c*_1_ is significantly worse. It achieves sparsity in just one of the two sequence datasets being compared, which is appropriate for comparing a huge dataset to a moderate-size dataset, e.g. many DNA reads to a moderate-size genome. It *might* be appropriate for comparing DNA reads to a human genome.

Sparsity in both datasets, with minimizers or word-restricted seeds, is appropriate for “huge-versus-huge” comparisons. A typical example is aligning DNA reads to each other in order to assemble them, which was a major motivation for minimizers [RHH^+^04]. Other examples are searching DNA sequences from unknown organisms against a multi-genome database, or checking if DNA data has contamination from other organisms [SS20].

### 4.3 Words versus minimizers

This study indicates that seeding at minimally-overlapping words is superior to minimizers. One caveat – bias due to reduced minimizer density at sequence edges – is addressed in the Supplement, and does not change this conclusion. It is important to note, however, that minimizer schemes are still being optimized [MPB^+^17, MDK18]. On the other hand, we have barely begun to optimize word-restricted seeding.

Compared to word-restricted seeds, a minimizer seed match has an extra contextual requirement. A seed match can be destroyed by a mutation inside the seed: this applies equally to both methods. However, minimizers experience an additional effect: a mutation outside the seed can make that seed position no longer a minimizer. This reduces the sensitivity of minimizers, but increases their specificity, which fits our observations.

Our word-restricted seeding has a potential disadvantage: there is no upper bound on distance between words. The probability of longer distance decreases *exponentially* in complex sequence, but not in simple sequence such as polypurine tracts or short-period tandem repeats. Pure simple-sequence similarities are typically not wanted, because their significance is hard to assess and they do not reliably indicate homology.

### 4.4 Further advantages of words

Word-restricted seeding has further advantages over minimizers. Firstly, it can be co-designed with subset seeds. Secondly, it seems likely that word positions can be found faster than minimizer positions. Thirdly, word-restricted seeding is more conducive to efficient indexes. Seed matches are usually found with an index data-structure. There are various kinds of index, but they often include a lookup table for any possible DNA sequence of some length *d*. This table can be reduced (or *d* increased) with word-restricted seeding, because only a subset of length-d words are ever considered.

### 4.5 Co-designed seed patterns

The sensitivity benefit of spaced and subset seeds can be enhanced by using, instead of one seed pattern, several co-designed patterns [BKS03, SB06]. Each pattern tends to find similarities that tend to be missed by the other patterns. This idea could be combined with word-restricted seeding. For example, we could use four different patterns, each starting with one of the minimally-overlapping words RRRY, RYRR, RYYR, YYYR (Table 3). Most interestingly, the best set of words may then not be minimally-overlapping ones, but rather words whose overlaps complement the seed patterns.

### 4.6 Open questions

Our study provides a new motivation for the problem of maximizing the number of non-overlapping words. For our purposes, minimally-overlapping words are especially useful, but we remain unsure how best to quantify overlap. Another challenge is how to search a large number of possible word sets for one with low overlap. More generally, we would like to design a set of word-restricted subset-seed patterns.

A further difficulty is how to optimize word-restricted seeding when the letter frequencies are unequal. In this case, we cannot simply seek an optimal set of *n* length-*k* words, because the sparsity is not constant. It is notable, however, that most natural DNA has near-equal frequencies of `r` and `y`.

Going further in the direction of empirical data, it might be useful to optimize word-restricted seeding for a particular sequence set (e.g. a genome). Presumably, it is beneficial to use words that are anti-clumped while tending to avoid repetitive sequence. Minimally-overlapping words avoid some kinds of repeat, e.g. homopolymers. Previously, minimizer ordering was defined by frequency in a particular sequence set [CLJ^+^14].

Word-restricted seeding requires fast word-finding. Perhaps some word sets are conducive to fast detection, e.g. the words in the last row of Table 3 share a common prefix.

When we use increasingly long minimum-variance words, with fixed sparsity *n*, the sensitivity might approach that of every-*n*th seeding (Figure 2, 3). The seed count of every-*n*th seeding has zero variance: can the words achieve arbitrarily-low variance? If so, they become arbitrarily close to a universal *k*-mer hitting set. Perhaps optimized minimizers, minimally-overlapping words, and universal *k*-mer hitting sets will converge.

## 5 Acknowledgments

We are grateful to Paul Horton for suggesting seeds starting with `a`, and Shotaro Tadachi for investigating word-free tracts in human DNA. GK was partially funded by RFBR, project 20-07-00652, and joint RFBR and JSPS project 20-51-50007. LN was partially funded by ANR, ASTER project ANR-16-CE23-0001.

## Reference

[AT18] Meznah Almutairy and Eric Torng. Comparing fixed sampling with minimizer sampling when using *k*-mer indexes to find maximal exact matches. PLoS ONE, 13(2), 2018.

[BKS03] Jeremy Buhler, Uri Keich, and Yanni Sun. Designing seeds for similarity search in genomic DNA. In Proceedings of the seventh annual international conference on Research in computational molecular biology, pages 67–75, 2003.

[Bla15] Simon R. Blackburn. Non-overlapping codes. IEEE Transactions on Information Theory, 61(9):4890–4894, 2015.

[CLJ+14] Rayan Chikhi, Antoine Limasset, Shaun Jackman, Jared T. Simpson, and Paul Medvedev. On the representation of de Bruijn graphs. In International conference on Research in computational molecular biology, pages 35–55. Springer, 2014.

[Csu04] Miklos Csrnös. Performing local sim ilarity searches with variable length seeds. In Annual Symposium on Combinatorial Pattern Matching, pages 373–387. Springer, 2004.

[DKGDG15] Sebastian Deorowicz, Marek Kokot, Szymon Grabowski, and Agnieszka Debudaj-Grabysz. KMC 2: fast and resource-frugal *k*-mer counting. Bioinformatics, 31(10):1569–1576, May 2015.

[FN14] Martin C. Frith and Laurent Noé. Im proved search heuristics find 20 000 new alignments between human and mouse genomes. Nucleic acids research, 42(7):e59–e59, 2014.

[HLO+16] Lars Hahn, Chris-André Leimeister, Rachid Ounit, Stefano Lonardi, and Burkhard Morgenstern. rasbhari: Optimizing spaced seeds for database searching, read mapping and alignment-free sequence comparison. PLoS computational biology, 12(10):e1005107, 2016.

[II07] Lucian Ilie and Silvana Ilie. Multiple spaced seeds for homology search. Bioinformatics, 23(22):2969–2977, 2007.

[JKD+18] Chirag Jain, Sergey Koren, Alexander Dilthey, Adam M. Phillippy, and Srinivas Aluru. A fast adaptive algorithm for computing whole-genome homology maps. Bioinformatics, 34(17):i748–i756, September 2018.

[KJ07] Janez Konc and Duŝanka Janežic. An improved branch and bound algorithm for the maximum clique problem. MATCH Commun. Math. Comput. Chem., 58:569–590, 2007.

[KNR06] Gregory Kucherov, Laurent Noé, and Mikhail Roytberg. A unifying framework for seed sensitivity and its application to subset seeds. J Bioinform Comput Biol, 4(2):553–569, Apr 2006.

[KWS+11] Szymon M. Kiełbasa, Raymond Wan, Kengo Sato, Paul Horton, and Martin C. Frith. Adaptive seeds tame genomic sequence comparison. Genome research, 21(3):487–493, 2011.

[Li18] Heng Li. Minimap2: pairwise alignment for nucleotide sequences. Bioinformatics, 34(18):3094–3100, September 2018.

[LKH+13] Yang Li, Pegah Kamousi, Fangqiu Han, Shengqi Yang, Xifeng Yan, and Subhash Suri. Memory efficient minimum substring partitioning. Proceedings of the VLDB Endowment, 6(3):169–180, January 2013.

[MDK18] Guillaume Marçais, Dan DeBlasio, and Carl Kingsford. Asymptotically optimal minimizers schemes. Bioinformatics, 34(13):i13–i22, 2018.

[MPB+17] Guillaume Marçais, David Pellow, Daniel Bork, Yaron Orenstein, Ron Shamir, and Carl Kingsford. Improving the performance of minimizers and winnowing schemes. Bioinformatics, 33(14):i110–i117, 2017.

[MTL02] Bin Ma, John Tromp, and Ming Li. PatternHunter: faster and more sensitive homology search. Bioinformatics, 18(3):440–445, Mar 2002.

[NK04] Laurent Noé and Gregory Kucherov. Improved hit criteria for DNA local alignment. BMC bioinformatics, 5(1):149, 2004.

[OPM+17] Yaron Orenstein, David Pellow, Guillaume Marçais, Ron Shamir, and Carl Kingsford. Designing small universal *k*-mer hitting sets for improved analysis of high-throughput sequencing. PLoS computational biology, 13(10):e1005777, 2017.

[RHH+04] Michael Roberts, Wayne Hayes, Brian R. Hunt, Stephen. M. Mount, and James A. Yorke. Reducing storage requirements for biological sequence comparison. Bioinformatics, 20(18):3363–3369, Dec 2004.

[SB06] Yanni Sun and Jeremy Buhler. Choosing the best heuristic for seeded alignment of DNA sequences. BMC bioinformatics, 7(1):133, 2006.

[SS20] Martin Steinegger and Steven L. Salzberg. Terminating contamination: large-scale search identifies more than 2,000,000 contaminated entries in GenBank. Genome biology, 21(1):1–12, 2020.

[SWA03] Saul Schleimer, Daniel S. Wilkerson, and Alex Aiken. Winnowing: local algorithms for document finger-printing. In Proceedings of the 2003 ACM SIGMOD international conference on Management of data, pages 76–85. ACM, 2003.

[Tam92] Koichiro Tamura. Estimation of the number of nucleotide substitutions when there are strong transition-transversion and G+C-content biases. Mol Biol Evol, 9(4):678–687, 1992.

[WS14] Derrick E. Wood and Steven L. Salzberg. Kraken: ultrafast metagenomic sequence classification using exact alignments. Genome Biol., 15(3):R46, 2014.

