## Supplement for "Minimally-overlapping words for sequence similarity search"

2020-08-17

#### 1 Every $n$th sparsity

Every-$n$th sparsity means to only use seeds starting at every $n$th position in one of the two sequences being compared. We considered two slightly different ways to do that: every $n$th position starting from the first position in the sequence, versus every $n$th position starting from one of the first $n$ positions chosen randomly. With the random-origin method, the average fraction of selected seed positions is precisely $1/n$. With the origin-1 method, on the other hand, the number of seed positions is the smallest integer such that the fraction is $\geq 1/n$.

Since we test sensitivity with fairly short length-100 sequences, the difference is not negligible (Figure S1). In the rest of this study, the random-origin method is used.

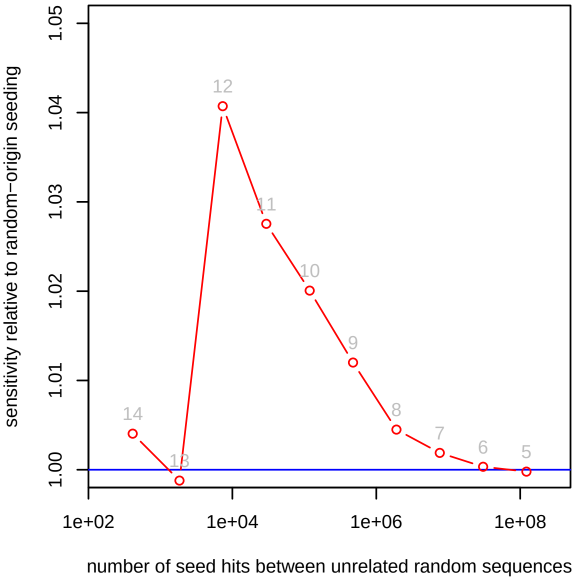


**Figure S1:** Sensitivity (y-axis) and spurious hit count (x-axis) for exact-match seeds with every-8th sparsity. Red: origin-1 seeding. Blue: random-origin seeding. Sensitivity was measured on sequence pairs with PAM distance 20. Seed lengths 5–14 were tested, shown in gray.

#### 2 Variance in count of words

In the main text, we derived an equation for the variance in number of occurrences of a set of words, in a random sequence of length $s$. That derivation omits some tedious algebra, so it may be hard to follow. Here is one intermediate step. Assuming that $s \geq 2k - 2$:

$$\begin{aligned} \mathrm{E}[X^2] = {} & (s-k+1)p \\ & + 2\sum_{l=1}^{k-1} \left[(s-k+1-l) \sum_{V,W} B_{VW}^{k-l} P_V^l P_W^k\right] \\ & + (s-2k+1)(s-2k+2)p^2 \,. \end{aligned}$$

Note that, if $s < 2k$, there is no pair of non-overlapping word positions, so the $p^2$ term should be 0. Thus, the above equation is correct when $s \geq 2k - 2$.

##### 2.1 Circular sequences

For a random circular sequence of length $s$,

$$\begin{aligned} \mathrm{E}[X^2] &= \mathrm{E}[(I_1 + I_2 + \cdots + I_s)^2] \\ &= \textstyle\sum_{i,j} \mathrm{E}[I_i I_j] \,. \end{aligned}$$

Let us assume that $s \geq 2k - 1$, to avoid cases where the end of word $V$ overlaps the start of word $W$ *and* the end of word $W$ overlaps the start of word $V$. If we define the modular distance $l =$

$\min(|i-j|,\, s-|i-j|)$, then

$$\mathrm{E}[I_i I_j] = \begin{cases} p & \text{if } l = 0 \\ \sum_{V,W} B_{VW}^{k-l} P_V^l P_W^k & \text{if } 0 < l < k \\ p^2 & \text{if } l \geq k\,. \end{cases}$$

Therefore

$$\begin{aligned}\mathrm{E}[X^2] = {} & sp \\ & + 2\sum_{l=1}^{k-1}\big[s\sum_{V,W} B_{VW}^{k-l} P_V^l P_W^k\big] \\ & + (s-2k+1)sp^2\,.\end{aligned}$$

### 3 Finding low-variance words

#### 3.1 Overlap scores

As a minor simplification, we minimized an integer "overlap score", which is equivalent to minimizing variance. This is possible because we only considered the case of equal letter frequencies. So $P_W^n$ (the product of probabilities of the first $n$ letters in word $W$) equals $a^{-n}$, where $a$ is the alphabet size (either 2 for the `ry` alphabet or 4 for the `acgt` alphabet). Thus, we can define an integer score:

$$\text{score} = \sum_{V,W}\sum_{l=1}^{k-1}[(s-k+1-l)B_{VW}^{k-l} a^{k-l-1}]\,,$$

such that lower score implies lower variance. For VMR1, the overlap score is simpler:

$$\text{score} = \sum_{V,W}\sum_{l=1}^{k-1}[B_{VW}^{l} a^{l-1}]\,.$$

#### 3.2 Simulated annealing

For simulated annealing, we began with a random set of words. Then, at each step of the algorithm, we proposed replacing a random word in the set with a random word not in the set. If this replacement would decrease the overlap score, it was accepted. If it would increase the overlap score, it was accepted with probability: $\exp[(\text{oldScore}-\text{newScore})/t]$. The temperature parameter $t$ was set equal to the overlap score of the initial random word set, then multiplied by 0.9999999 at each iteration. The algorithm was halted when $t \leq 0.05$ (because the probability of accepting any integer score increase is extremely low then).

**Table S1:** Variance-to-mean ratios.

| words | VMR1 | VMR2 |
|---|---|---|
| *sparsity 4* | | |
| `ryn` | 0.25 | 0.417 |
| `atc,atg,caa,cac,cag,ccg,ctc,ctg,gaa,gcg,gtg,taa,tac,tag,ttc,ttg` | 0.219 | 0.406 |
| `rrrrryn,rryrryn,rryryyn,ryrrrrn,ryrrryn,ryryryn,ryyrrrn,ryyrryn,ryyryrn,ryyryyn,ryyyryn,ryyyyyn,yryrryn,yyyrrrn,yyyrryn,yyyyryn` | 0.0938 | 0.174 |
| `rrryrrr,rrryrry,rrryryy,rrryyrr,rrryyry,rrryyyr,rryryrr,rryryry,rryyyyr,rryyyyy,yrrrrrr,yrrrrry,yrryrrr,yrryrry,yrryryy,yrryyrr,yrryyry,yrryyyr,yryryrr,yryryry,yryyrrr,yryyrry,yryyyrr,yryyyry,yryyyyr,yryyyyy,yyrrrry,yyryrrr,yyryrry,yyryryy,yyryyry,yyyyyyr` | 0.0898 | 0.172 |

### 4 Low-variance words

In the main text, we saw that sensitivity of word-restricted seeding can be boosted by using words with low variance (VMR1 and VMR2). Some further, less impressive, results are shown here.

Firstly, we sought even longer and lower-variance words, by heuristic search (simulated annealing). We found a set of 32 length-7 words (Table S1), which have lower VMR1 and VMR2 than the sixteen length-6 words. Surprisingly, however, their sensitivity is slightly worse (Figure S2). This indicates that neither VMR1 nor VMR2 is a perfect guide to sensitivity.

Secondly, we sought low-variance DNA words without restriction to a 2-letter alphabet, by heuristic search (simulated annealing). In particular, we sought to replace `ryn` with a less-overlapping set of length-3 words. We found a set with lower VMR1 and VMR2 (Table S1: `atc`, `atg`, `caa`, ...). The sensitivity of seeds starting at these words is indeed better than that of seeds starting at `ry` (Figure S2).

### 5 Edge bias of minimizers

In the main text, we found minimizer positions by sliding a length-$w$ window along the sequence, and getting all positions that are the minimum of any

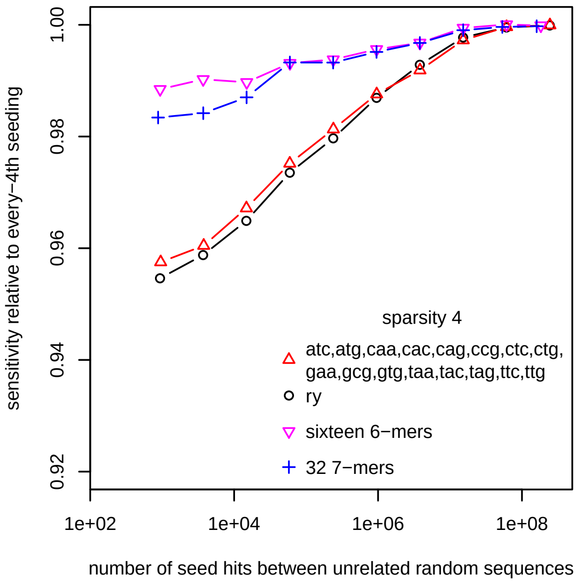


**Figure S2:** Sensitivity (y-axis) and spurious hit count (x-axis) for exact-match seeds with word-based sparsity. Sensitivity was measured on sequence pairs with PAM distance 20. Seed lengths 5–14 were tested.

window. This procedure, termed "interior minimizers" [RHH+04], means that positions near sequence edges have fewer chances to be a minimizer. In particular, the first position in a sequence has only one chance to be a minimizer, because it is contained in only one length-$w$ window. Thus, minimizer density is lower, on average, near sequence edges.

Moreover, in the main text we measured minimizer density in length-$10^6$ sequences, but sensitivity was tested with length-100 sequences. This makes our tests biased against minimizers, because their density in the tested length-100 sequences is lower than their reported density in length-$10^6$ sequences, due to the edge effect.

To address this concern, we performed a further test of minimizer sensitivity with length-100 sequences. This time, when finding minimizer positions, we treated the sequences as circular, so there is no density reduction at sequence edges.

Figure 6 in the main text, when modified by using "circular minimizers", becomes Figure S3 here. Recall that Figure S3A,B compares words to minimizers with slightly lower sparsity (higher density), giving an unfair advantage to the minimizers. As expected, circular minimizers are slightly more sensitive than interior minimizers (Figure S3 versus Figure 6). The main conclusion does not change: minimally-overlapping words perform better than minimizers.

### References

[RHH+04] Michael Roberts, Wayne Hayes, Brian R. Hunt, Stephen. M. Mount, and James A. Yorke. Reducing storage requirements for biological sequence comparison. *Bioinformatics*, 20(18):3363–3369, Dec 2004.

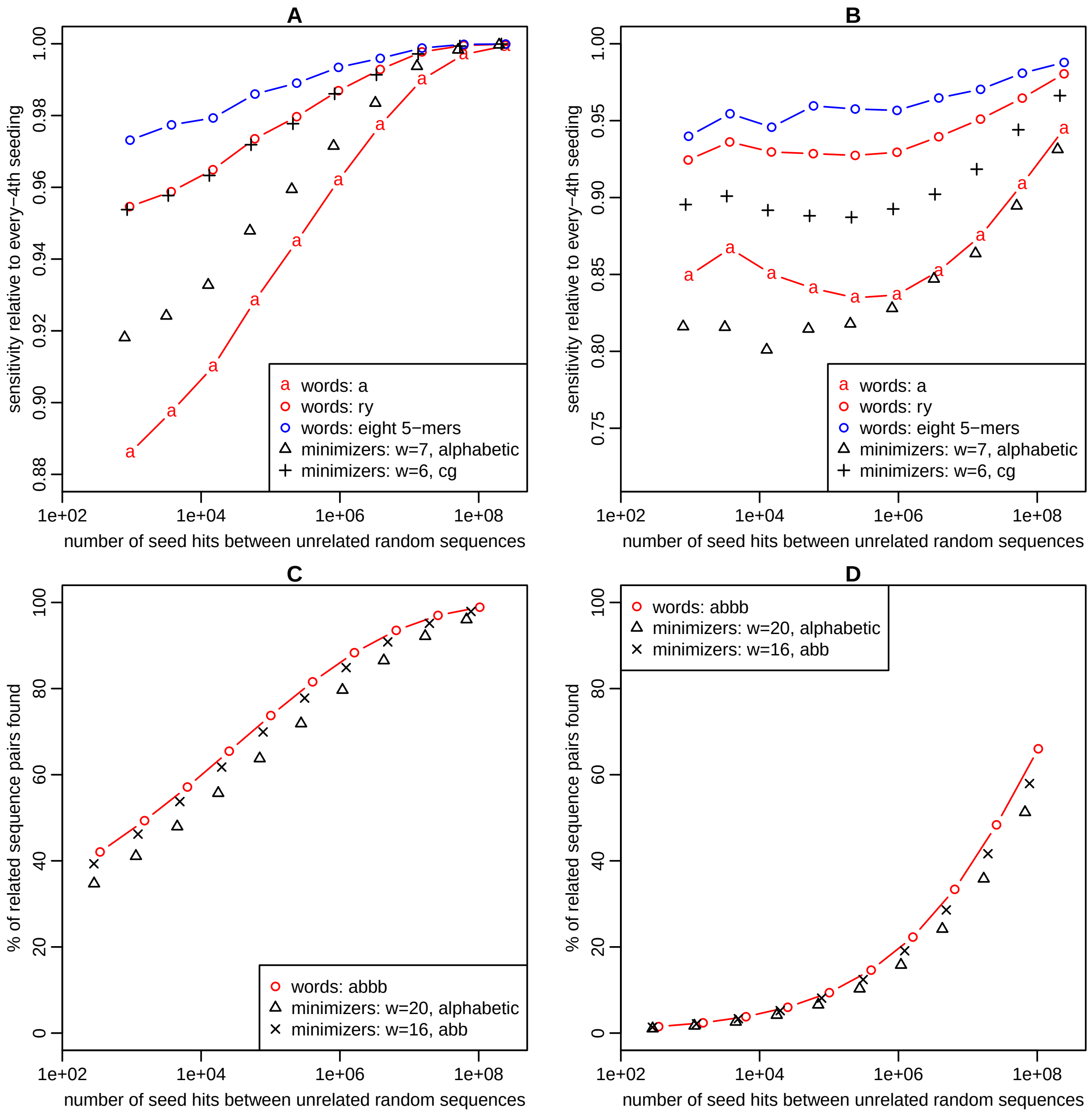


**Figure S3:** Sensitivity (y-axis) and spurious hit count (x-axis) for exact-match seeds at either word positions or minimizer positions. Seed lengths 5–14 were tested. Sensitivity was measured on sequence pairs with PAM distance 20 (**A**, **C**) or 50 (**B**, **D**). Here, "circular minimizers" were used.
